# Attractor-like circuits improve visual decoding and behavior in zebrafish

**DOI:** 10.1101/2024.02.03.578596

**Authors:** Martin Privat, Enrique Carlos Arnoldo Hansen, Thomas Pietri, Emiliano Marachlian, Alejandro Uribe-Arias, Auriane Duchemin, Virginie Candat, Sarah Nourin, Germán Sumbre

**Affiliations:** Institut de Biologie de l’ENS (IBENS), Département de biologie, École normale supérieure, CNRS, INSERM, Université PSL, 75005 Paris, France

## Abstract

Attractor networks are neural circuits with stable states that represent information or memories. They play a crucial role in memory retrieval, decision-making and integrating noisy cues. In zebrafish larvae, the spontaneous dynamics of the optic tectum is structured according to topographically organized neuronal assemblies exhibiting attractor-like behavior.

Here, we took advantage of the Methyl-CpG-binding protein 2 (MeCP2) deficient zebrafish mutant, which displays perturbed tectal dynamics, to study the functional role of the attractor-like circuits in visual processing.

In comparison to wild-type larvae, the *mecp2*-mutant showed reduced functional connectivity in the optic tectum. This abnormal connectivity significantly affected the visual response, and the ability to discriminate between visual stimuli. Finally, the mutant larvae where less efficient in hunting paramecia. We argue that the attractor dynamics of the tectal assemblies improve stimulus discrimination, visual resolution, and increase the sensitivity to behaviorally relevant visual stimuli.

## INTRODUCTION

Across brain regions, neuronal circuits display different structural architectures, dynamics and functional roles, usually organized according to neuronal assemblies (a group of highly correlated neurons). Certain neuronal circuits show activity patterns with attractor-like dynamics, based on recurrent connectivity. These attractor-like circuits display temporal dynamics that settle into self-sustained stable activity patterns. This ability to converge and maintain specific activity states allows these circuits to recall stored patterns from partial cues (*i.e.*, pattern completion) or to discriminate and categorize its inputs (*i.e.*, pattern separation). Such attractor-like circuits were used to model short-term memory ^1–3^, to explain head-direction cells and eye fixations ^4–9^, and as a mechanism for the representation of head direction in drosophila and orientation tuning in primary visual cortex ^5,10,11^. Multi-dimensional continuum attractors have been suggested as models to explain place and grid cells ^12^ and proposed for faster odor recognition in the olfactory bulb ^13^. Therefore, neuronal attractors could reflect basic building blocks of the brain ^14,15^.

To learn about the contribution of attractor dynamics to the circuit’s functional role, it is necessary to disrupt *in* vivo the recurrent functional connectivity, while keeping intact the circuit’s inputs. However, to experimentally perturb the local connectivity without affecting the circuit’s afferents, is technically unfeasible.

In the zebrafish larva, the optic tectum plays a role in detecting the physical properties of visual stimuli, it processes them and generates goal-directed motor behaviors. The tectum’s ongoing spontaneous activity is organized according to topographically compact neuronal assemblies ^16–21^. These assemblies mimic visually induced responses associated with the larva’s prey, and are organized according to the tectum’s retinotopic map ^16,18^. Their spontaneous activation represent ‘‘preferred’’ network states and mutual inhibition, characteristics reminiscent of attractor-like circuits ^16^.

To understand the functional role of the attractor dynamics of these neuronal assemblies, we took advantage of the Methyl-CpG-binding protein 2 (MeCP2) deficient zebrafish mutant, that displays disrupted local functional connectivity in the optic tectum (significantly weaker pairwise correlations between the spontaneous activity of tectal neurons).

Even though homozygous null *mecp2* mutations in humans are embryonic lethal, the *mecp2*-mutant zebrafish larvae do not show any major morphological or behavioral phenotypes ^22^. Their retina functionally innervates the optic tectum, allowing the mutants to perform visually guided behaviors. Therefore, the mecp2^Q63*/Q63^* mutant (*mecp2*-mutant) zebrafish represents an ideal model to study the functional role of neuronal assemblies with attractor-like dynamics in visual processing. Using the *mecp2*-mutant larvae, we observed that the reduced functional connectivity (lack of neuronal assemblies) affected the circuit visual responses in the optic tectum. The *mecp2*-mutant larvae showed reduced amplitude, reliability, and duration of the visual responses, and poor decoding of the visual stimulus. However, when the visual stimulus was larger, the number of responsive neurons in the *mecp2*-mutant larvae was larger. These results match those expected from attractor-like circuits (*e.g.*. pattern completion, sustained activity, and pattern discrimination). Furthermore, the abnormal visual response was associated with perturbed prey-capture behavior. These results suggest that the intrinsic functional connectivity of the optic tectum improves visual spatial resolution allowing a better detection of prey-like stimuli.

## RESULTS

### The optic tectum of *mecp2-*mutant larvae shows abnormal spontaneous activity

In sensory brain areas, the spontaneous neuronal activity displays a spatiotemporal structure that reflects the functional connectivity of neural circuits ^23–25^. To evaluate the functional connectivity in the *mecp2*-mutant larvae, we studied the spontaneous activity of the optic tectum in both *mecp2*-mutant and wild-type larvae. To that end, we combined two-photon microscopy and transgenic wild-type and mutant larvae (5 dpf), expressing pan-neuronally, the genetically encoded Ca^2+^ indicator, GCaMP3 (Fig. 1A). Using this approach, we monitored the neuronal activity of the periventricular neurons (PVN) for 35 min in absence of sensory stimulation, at a frame-rate of ∼4 Hz, in paralyzed larvae (wild type: 721.33 ± 59 neurons / larva, n=9 and *mecp2*-mutant : 721.6 ± 37 neurons / larva, n=10).

**Figure 1:**
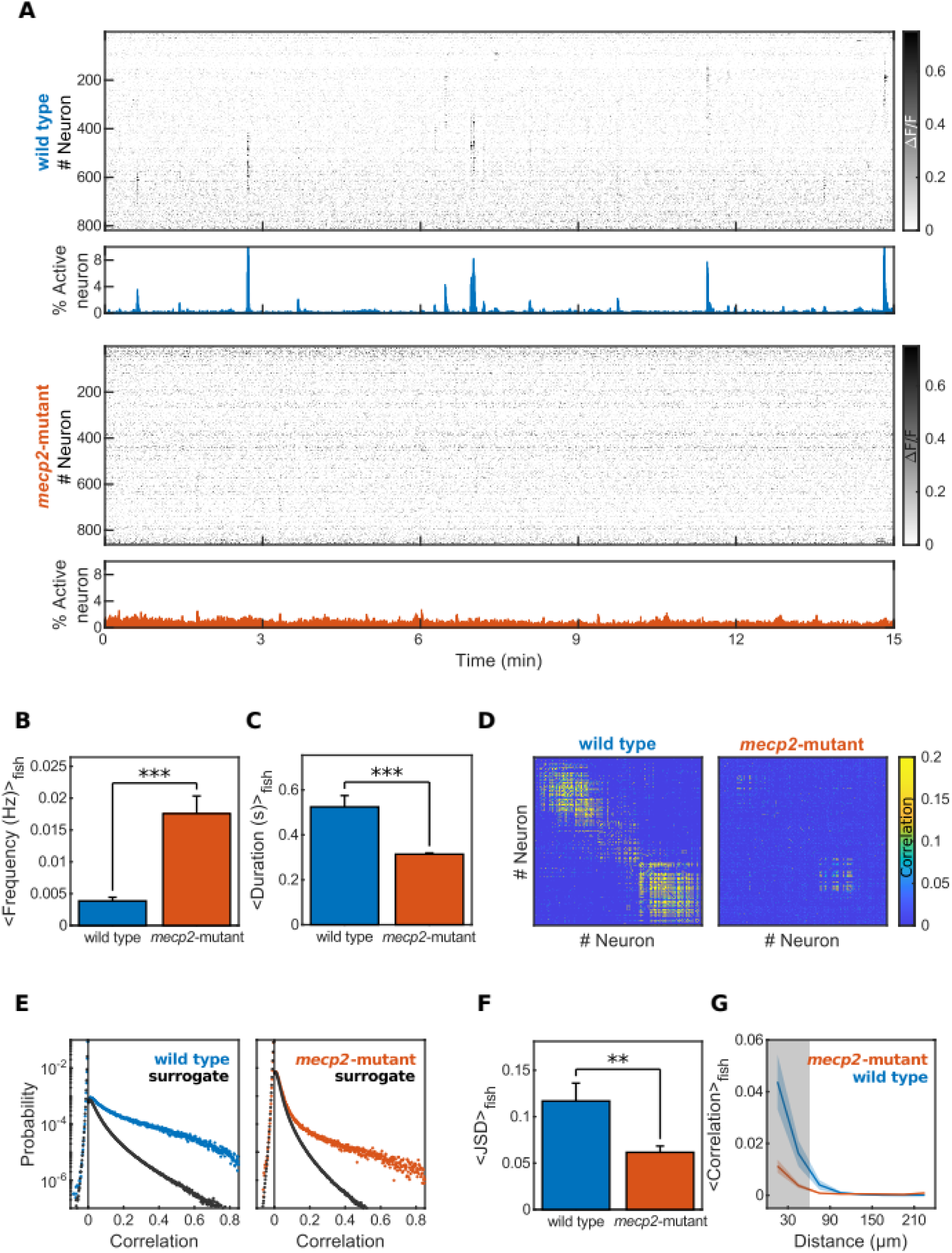
The optic tectum of the mecp2-mutant larva display reduced functional connectivity. (A) A representative raster plot of the spontaneous activity in the optic tectum of wild-type (top) and *mecp2-*mutant (bottom) larvae (5 dpf). Note the synchronous events in the wild-type larva (top) compared to the non-synchronous events in the *mecp2*-mutant larva (bottom). Gray scale: relative variation of fluorescence (ΔF/F). (B) Average frequency of the Ca^2+^-transients in wild-type (blue, 3.8 ± 0.6 x10^-3^ Hz, n=9) and *mecp2-*mutant (orange, 17.6 ± 2.8 x10^-3^ Hz, n=10) larvae, p=4.33e-5 (bilateral Wilcoxon rank sum test). Error bars: standard error. (C) Average duration of the Ca^2+^-transients in wild-type (blue, 524 ± 50 ms, n=9) and *mecp2-*mutant (orange, 314 ± 6 ms, n=10) larvae, p=2.16e-5 (bilateral Wilcoxon rank sum test). Error bars: standard error. (D) Example of pairwise Pearson’s correlation between the spontaneous activity of tectal neurons in 5 dpf wild-type and *mecp2-*mutant larvae. Colorbar: correlation values. (E) Distribution of the pairwise correlations between the spontaneous activity of tectal neurons in wild-type (blue) and *mecp2-*mutant (orange) larvae, and their respective surrogate data sets (black). (F) The average distance between the distributions of the pairwise correlation coefficients and their respective surrogates in wild-type (blue, 0.12 ± 0.02, n=9) and *mecp2-*mutant (orange, 0.06 ± 0.006, n=10) larvae, quantified as Jensen-Shannon distance (JSD), p=5.67e-3 (bilateral Wilcoxon rank sum test). Error bars: standard error. (G) The average relationship between the physical distance of each pair of neurons and their coefficient of correlation in wild-type (blue, n=9) and *mecp2-*mutant (orange, n=10) larvae. The data have been pooled in 50 μm bins. Gray area: p<0.05 (bilateral Wilcoxon rank sum test).

We found that the ongoing spontaneous activity in the optic tectum of *mecp2*-mutant larvae was different from that of wild-type larvae. We observed that the frequency of spontaneous Ca^2+^ transients in *mecp2*-mutant larvae was significantly higher than in wild types (wild type : 17.6 ± 2.8 x10^-3^ Hz, n=9 versus *mecp2*-mutant: 3.8 ± 0.6 x10^-3^ Hz, n=10; bilateral ranksum test, p=4.33e-05; Fig. 1B). The duration of their Ca^2+^-events was significantly shorter in *mecp2*-mutant with respect to that of wild-type larvae (wild type: 524 ± 50 ms, n=9 versus *mecp2*-mutant: 314 ± 6 ms, n=10; bilateral ranksum test, p=2.165e-05; Fig. 1C). In addition, the Pearson’s pairwise correlations between the spontaneous activity of tectal neurons in the *mecp2*-mutant were significantly lower than in wild-type larvae (Fig. 1D). To take into account the possible influence of the level of activity on the measure of the correlation coefficients (the level of activity of the *mecp2*-mutant tectal circuit were more than seven fold that of the wild-type larvae), we computed the Jensen-Shannon distance (JSD) between the distribution of pairwise correlation coefficients between neurons (Fig. 1E), and that of a null model, taking into account the baseline activity of each tectal neuron (see Methods). Briefly, the JSD measures the difference between two distributions: a small JSD (close to zero) indicates that the distributions are very similar, while a large JSD indicates that the distributions differ. In our case the JSD is used to quantify the pairwise correlations with respect to the null-model (see methods). In the optic tectum of *mecp2*-mutant, the spontaneous activity exhibited a significantly lower JSD than that of wild-type larvae (wild type: 0.12 ± 0.02, n=9; *mecp2*-mutant: 0.06 ± 0.006, n=10, bilateral ranksum test, p=5.67e-3; Fig. 1E and 1F), demonstrating a lower level of correlations between neurons. These results suggest that *mecp2*-mutant larvae show abnormal functional connectivity between tectal neurons.

We then analyzed the relationship between the spontaneous activity correlations and the Euclidean distance between neurons. We found that at short distances (< 60 μm), *mecp2*-mutant larvae showed significantly lower correlations (Fig. 1G).

The *mecp2* mutation can lead to a bias in the generation or differentiation of neurons affecting the exctitatory/inhibitory ratio ^26–29^. Therefore, to investigate whether the abnormal spontaneous activity in the optic tectum is a result of an unbalanced neuronal circuit, we quantified the glutamatergic and GABAergic neuronal population in the optic tectum (Fig. S1,A). For this, we used double transgenic larvae expressing red and green fluorescence proteins under the vglut2a (excitatory) and gadb1 (inhibitory) promoters (see Methods). We did not find any significant difference in the number of both populations of neurons between the wild-type and *mecp2*-mutant larvae (Fig. S1,B-C), nor in their spatial distribution across the optic tectum (Fig. S1,D-F).

In the zebrafish optic tectum, the ongoing spontaneous activity is organized according to neuronal assemblies composed of highly correlated neurons. These neuronal assemblies are spatially organized reflecting the functional retinotopic map, they are tuned to biologically relevant visual stimuli (e.g., prey), and they show attractor-like dynamics ^16,18,19^. We therefore took advantage of the *mecp2*-mutant which shows a disrupted spatiotemporal organization (lack of neuronal assemblies) of the tectal spontaneous activity as a model to study the functional role of the neuronal assemblies in the zebrafish optic tectum.

### *mecp2*-mutant larvae show abnormal visual responses

To study the functional role of the tectal neuronal assemblies in visual processing, we first presented to *mecp2*-mutant and wild-type GCaMP3 larvae visual stimuli of different contrasts, while monitoring the neuronal response. The visual stimuli consisted of a looming circle (150 ms to full expansion) of a size of 40° centered at 30° to the frontal (0°) azimuth. The stimuli were presented at different contrasts (from 0.1 to 0.8 of the projectors contrast range), and repeated 15 times. To avoid habituation, the stimuli were presented in a pseudo-random order. In average, we recorded 612.29 ± 11.27 neurons in wild-type larvae (n=7), and 552 ± 5.27 neurons in *mecp2*-mutant larvae (n=10). To analyze the visually evoked responses in the tectal circuit, we first identified the neurons that significantly responded to the stimuli (see methods and ^16^). Both wild-type and *mecp2*-mutant larvae showed a larger number of responsive neurons as the contrast of the stimulus increased (Fig. 2A). The proportion of responsive neurons was significantly larger in the *mecp2*-mutant larvae with respect to the wild type (two-way anova, p=7.63e-14 for *mecp2-*mutant vs wild type and p=1.89e-04 for stimulus contrast, Fig. 2B). In contrast, the integral of the Ca^2+^ response (two-way anova using one level for *mecp2*-mutant/wild type and one level for stimulus contrast, p=1.32e-11 for *mecp2*-mutant vs. wild type, and p=1.40e-11 for contrast). Post-hoc bilateral ranksum tests were performed to locate significant differences (Fig. 2C). The peak response probability that describes how reliable the response is across trials was significantly higher in wild type in comparisson to *mecp2*-mutant larvae (two-way anova, d.f.=1, F=112.81, p=3.58e-19 for *mecp2*-mutant vs wild type and d.f.=7, F=5.81, p=7.53e-06 for stimulus contrast, Fig. 2D). The duration of the Ca^2+^ response was also longer in wild-type vs. *mecp2*-mutant larvae (two-way anova, p=1.50e-05 for *mecp2*-mutant vs. wild type and p=1.24e-05 for stimulus contrast, Fig. 2E).

**Figure 2:**
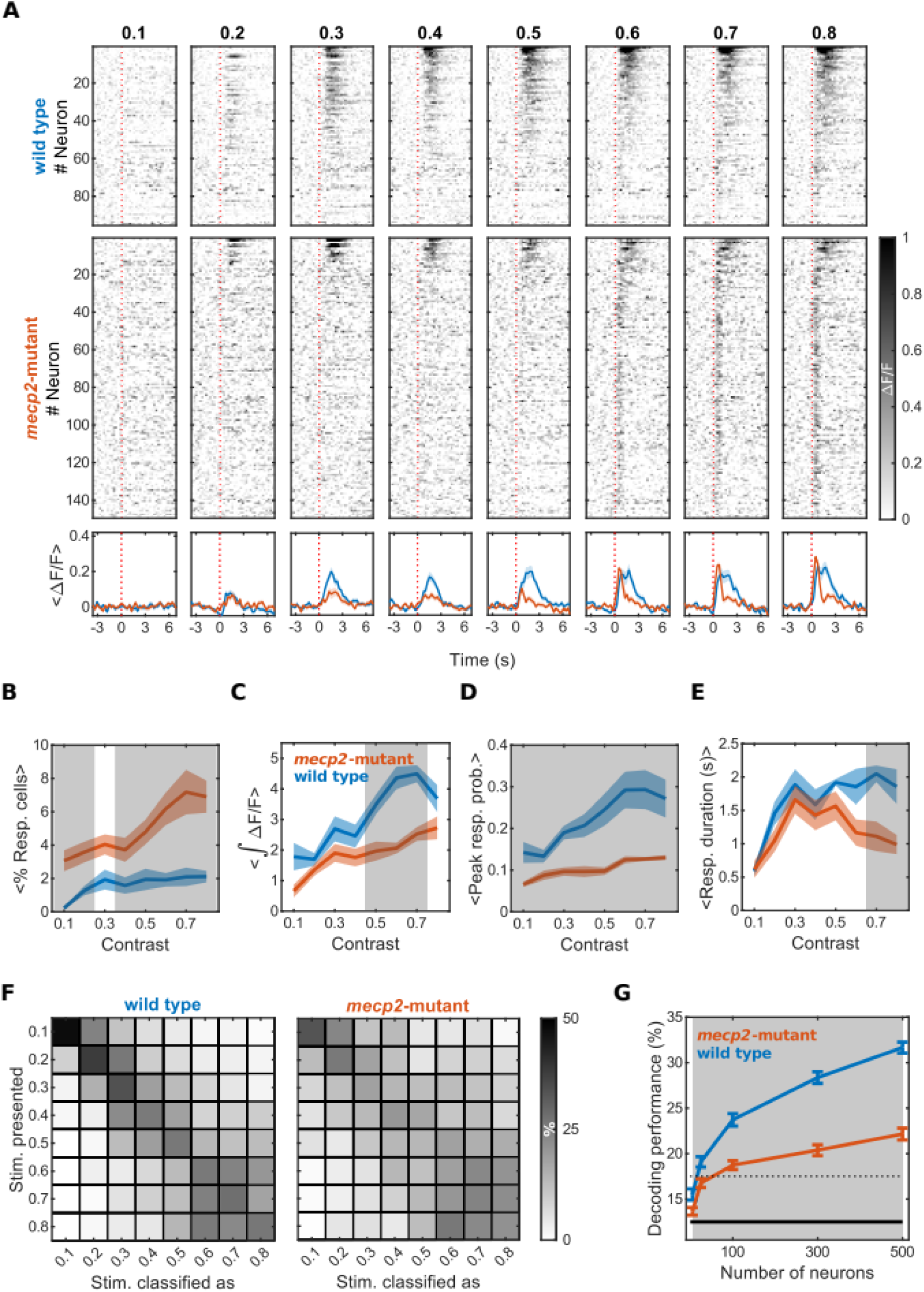
The optic tectum of the mecp2-mutant larva display abnormal responses to large visual stimuli. (A) Representative raster plots of the average induced activity across trials for different light contrasts. Top: wild-type larva. Middle: *mecp2-*mutant (middle panel) larva. Bottom: Average ± standard error of the wild-type (blue) and *mecp2-*mutant (orange) larvae. The red vertical dashed line represents the onset of light stimulation. (B) The averaged integral of the visually induced response for the different light contrasts across all wild-type (blue), or *mecp2-*mutant (orange) larvae. Shaded area: standard error. Gray area: p<0.05 (bilateral Wilcoxon rank sum test). (C) The averaged response probability for the different light contrasts across all wild-type (blue), or *mecp2-*mutant (orange) larvae. Shaded area: standard error. Gray area: p<0.05 (bilateral Wilcoxon rank sum test). (D) The average duration (full width at half maximum) of the visually induced response for all light contrasts across all wild-type (blue), or *mecp2-*mutant (orange) larvae. Shaded area: standard error. Gray area: p<0.05 (bilateral Wilcoxon rank sum test). (E) The averaged proportion of neurons responsive to the the different light contrasts across all wild-type (blue), or *mecp2-*mutant (orange) larvae. Note that in contrast to all other measurements (B, C, D), *mecp2-*mutant larvae show a significantly higher proportion of responsive neurons to the different light contrasts. Gray area: p<0.05 (bilateral Wilcoxon rank sum test). Shaded area: standard error. Gray area: p<0.05 (bilateral Wilcoxon rank sum test). (F) The confusion matrices resulting from the decoding of the light intensity presented to the wild-type (left panel) and *mecp2-*mutant (right panel) larvae. The percentage of correct (in the diagonal) or wrong decoding (outside the diagonal) is color-coded according to the scale bar. (G) The mean decoder performance (accuracy, *i.e.* proportion of correctly classified trials) for all wild-type (blue) and *mecp2-*mutant (orange) larvae. Horizontal solid bar: chance level, 1/(number of stimulus type). Horizontal dashed line: 95^th^ percentile of a binomial distribution. Error bars: standard error. Gray area: p<0.05 (paired bilateral t-test).

Overall, these results suggest that tectal visual responses in the the tectum of wild-type larvae are stronger, more reliable, more sustained and more spatially limited (smaller proportion of neurons) than in the *mecp2*-mutant larvae.

To test whether the difference in the visually induced neuronal responses between wild-type and *mecp2*-mutant larvae affected the capacity of the optic tectum to represent contrast information, we a maximum-likelihood classifier (see Methods, Fig. 2F). In order to estimate the number of cells necessary to reliably encode the stimulus information, we selected random ensembles of 5, 25, 100, 300 and 500 neurons (10 repetitions for each ensemble size) among the recorded neurons.

For a population of 500 neurons, on average the maximum likelihood decoder resulted in a better classification performance (proportion of trials correctly classified) for the wild-type larvae than for the *mecp2*-mutant larvae (Fig. 2G, wild type: 30.08% ± 0.36%, n=7 larvae x 10 repetitions versus *mecp2*-mutant: 21.30% ± 0.36%, n=10 larvae x 10 repetitions, p<0.001 bilateral two-sample t-test, d.f.=168, t-score=10.22, chance level is 12.5%, 95^th^ percentile binomial distribution 17.5%).

Previous studies showed that the optic tectum is essential for prey-capture behavior ^30^. In addition, the neuronal assemblies in the optic tectum are tuned to the angular size and spatial position relevant for the detection of prey ^16^. Therefore, we studied how disrupted functional connectivity did affect the visual response to prey-like visual stimuli. For this purpose, we presented to larvae expressing pan-neuronally GCaMP5, small light dots (5° in size) at different positions in the larva’s field of view (15°, 35°, 40° and 50°), while monitoring the tectum’s visual responses using a two-photon microscope.

To compare the neuronal responses to the four different stimulus positions between wild-type and *mecp2*-mutant larvae, we first identified neurons that significantly responded to the visual stimuli using a bootstrapping methods (see Methods). We found that the neuronal response to the different stimuli involved larger Ca^2+^ transients (integral of the Ca^2+^ transients: wild type= 0.73 ± 0.095, n=7 versus *mecp2*-mutant= 0.41 ± 0.048, n=9; p=0.016, bilateral Wilcoxon ranksum test; Fig. 3B), and more reliable (wild type: 0.055 ± 0.0062, n=7 versus *mecp2*-mutant: 0.037 ± 0.004, p=0.042, n=9, bilateral Wilcoxon ranksum test; Fig. 3C) in the wild-type compared to the *mecp2-*mutant larvae. On the contrary to what we found for wide-field stimuli of different contrasts, the proportion of neurons responsive to small light-spot stimuli was larger in wild-type larvae than in *mecp2-*mutant larvae (wild type: 8.68 ± 0.89%, n=7 versus *mecp2-*mutant: 3.80 ± 0.79%, n=9, p=0.0021, bilateral Wilcoxon ranksum test; Fig. 3D).

**Figure 3:**
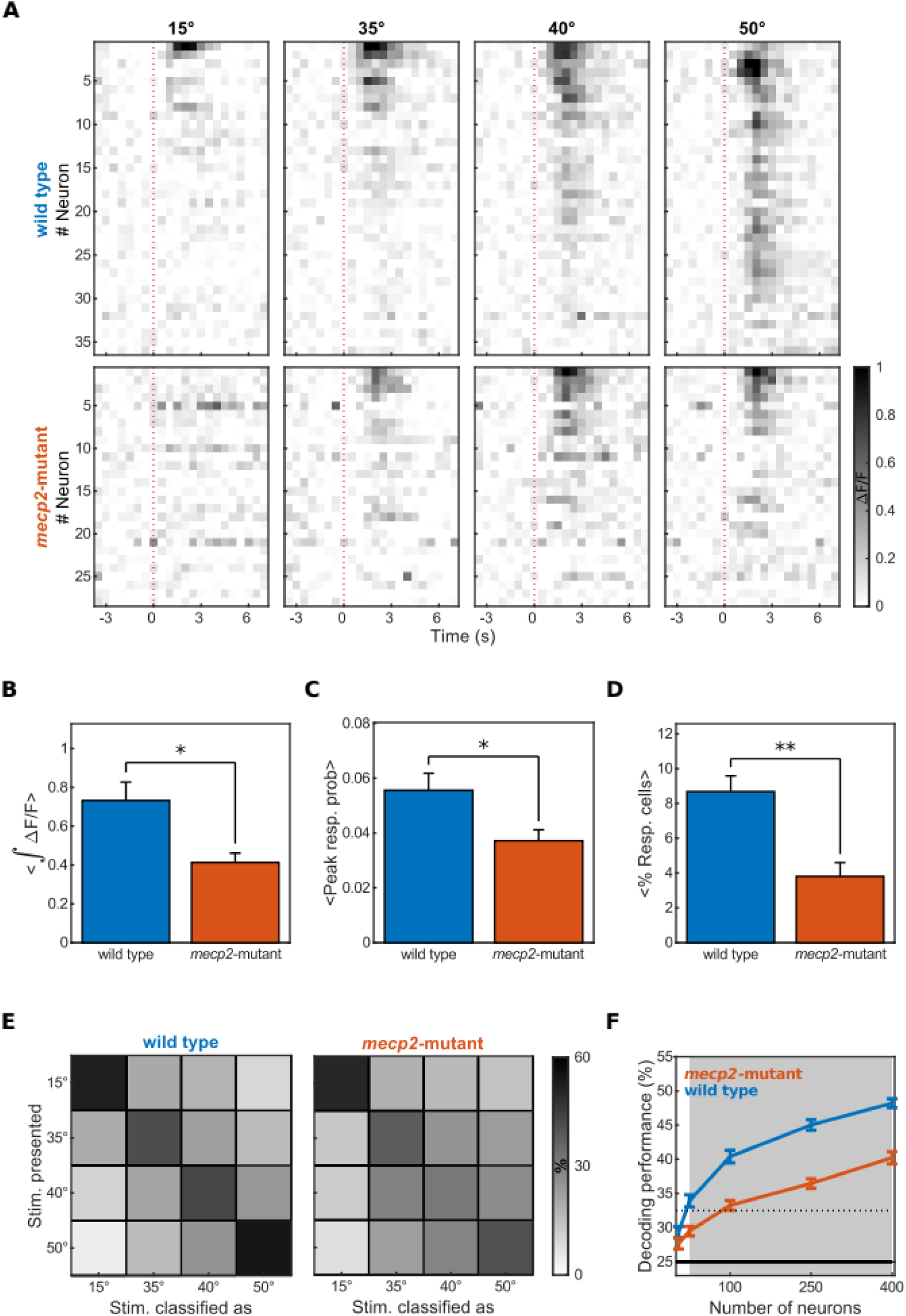
The optic tectum of the mecp2-mutant larva display abnormal responses to prey-like visual stimuli. (A) Representative raster plot of the average response to light-spot stimuli (5°), presented at 4 different azimuth angles (15°, 35°, 40° and 50° with respect to the caudo-rostral axis of the larva) in a wild-type (top) and *mecp2-*mutant larva (bottom). The red dashed line corresponds to the onset of the stimulus (t=0 s). (B) The average integral of the light-spots responses across all wild-type (blue, 0.73 ± 0.095, n=7), or *mecp2-*mutant (orange, 0.41 ± 0.048, n=9) larvae. p=1.64e-2, bilateral Wilcoxon ranksum test. Error bars: standard error. (C) The average peak response probability to the different light-spots across all wild-type (blue, 0.055 ± 0.0062, n=7), or *mecp2-*mutant (orange, 0.037 ± 0.004, n=9) larvae. p=4.17e-2, bilateral Wilcoxon ranksum test. Error bars: standard error. (D) The average proportion of responsive neurons to the light-spot stimuli across all wild-type (blue, 8.68 ± 0.89%, n=7), and *mecp2-*mutant (orange, 3.80 ± 0.79%, n=9) larvae. p=2.09e-3, bilateral Wilcoxon ranksum test. Error bars: standard error. (E) The average confusion matrices for a maximum-likelihood classifier trained using a leave-one-out approach, for 400 neurons selected at random in the optic tectum for wild-type (left, n=7) and *mecp2-*mutant larvae (right, n=9). Gray scale bar: the proportion of stimuli correctly classified. (F) The classifier performance (accuracy, *i.e.* proportion of correctly classified trials) for decoders on randomly selected pools of neurons of 5, 20, 50, 150, 300 and 400 neurons, for wild-type (blue, n=7) and *mecp2-*mutant (orange, n=9) larvae. Black horizontal solid line: chance level. Black horizontal dotted line: 95^th^ percentile for a binomial distribution. Shaded gray area: p<0.05 (paired bilateral t-test ). Error bars: standard error.

To test whether disrupted functional connectivity affects the capacity of the tectal circuit to better discriminate the location of prey-like visual stimuli (better spatial resolution), we classified the responses using a maximum likelihood decoder (MLD, see methods, Fig. 3E and Supp. Movie 1 ). We found that for a population of 400 neurons selected at random (10 repetitions), on average the maximum likelihood decoder resulted in a better classification performance (proportion of trials correctly classified) for wild-type larvae with respect to *mecp2-*mutant larvae (wild type: 48.2% ± 0.66%, n=7 larvae, *mecp2-*mutant: 40.22% ± 0.88%, n=9 larvae; p<0.001 bilateral two-sample t-test, chance level is 25%, 95^th^ percentile binomial distribution 32.5%, Fig. 3F).

### *mecp2-*mutant larvae show mild impairments during prey capture

In zebrafish larva, the optic tectum plays an important role in prey-capture behavior ^30^. Therefore, we tested whether the impaired decoding of visual stimuli, due to the lack of neuronal assemblies, did affect prey-capture behavior. For this purpose, we introduced in a recording chamber, 5 larvae and 24.8 ± 8.8 paramecia. We monitored for 30 min., the number of paramecia. The prey-capture assay was recorded at 100 Hz using a high-speed infrared camera placed above the recording chamber (Baumer TXG02). We observed that at the end of the 30 min, wild-type larvae ate more paramecia than the *mecp2-*mutant larvae (wild type: 4.06 ± 1.23, n=9 vs *mecp2-*mutant: 3.18 ± 1.70, n=9), but the difference was not significant (p=0.164, right-tailed Wilcoxon ranksum test), meaning that the *mecp2-*mutant larvae were still able to successfully capture paramecia (Fig. 4A). However, given the large number of paramecia used (24.8 ± 8.8), it is likely that larvae satiated before the end of the 30 min essay. It is therefore possible, that the *mecp2-*mutant larvae compensated a lower rate of capture by eating over a longer period of time. To test this hypothesis, we analyzed the first minutes of the recordings. Indeed, we found that *mecp2-*mutant larvae ate significantly less paramecia than wild-type larvae, during the early phase (1.5 to 4.5 min) of the assay (wild type: 2.28 ± 1.15, n=9, *mecp2-*mutant: 1.04 ± 0.58, n=9; p=0.015 right-tailed Wilcoxon ranksum test, fig 4A).

**Figure 4:**
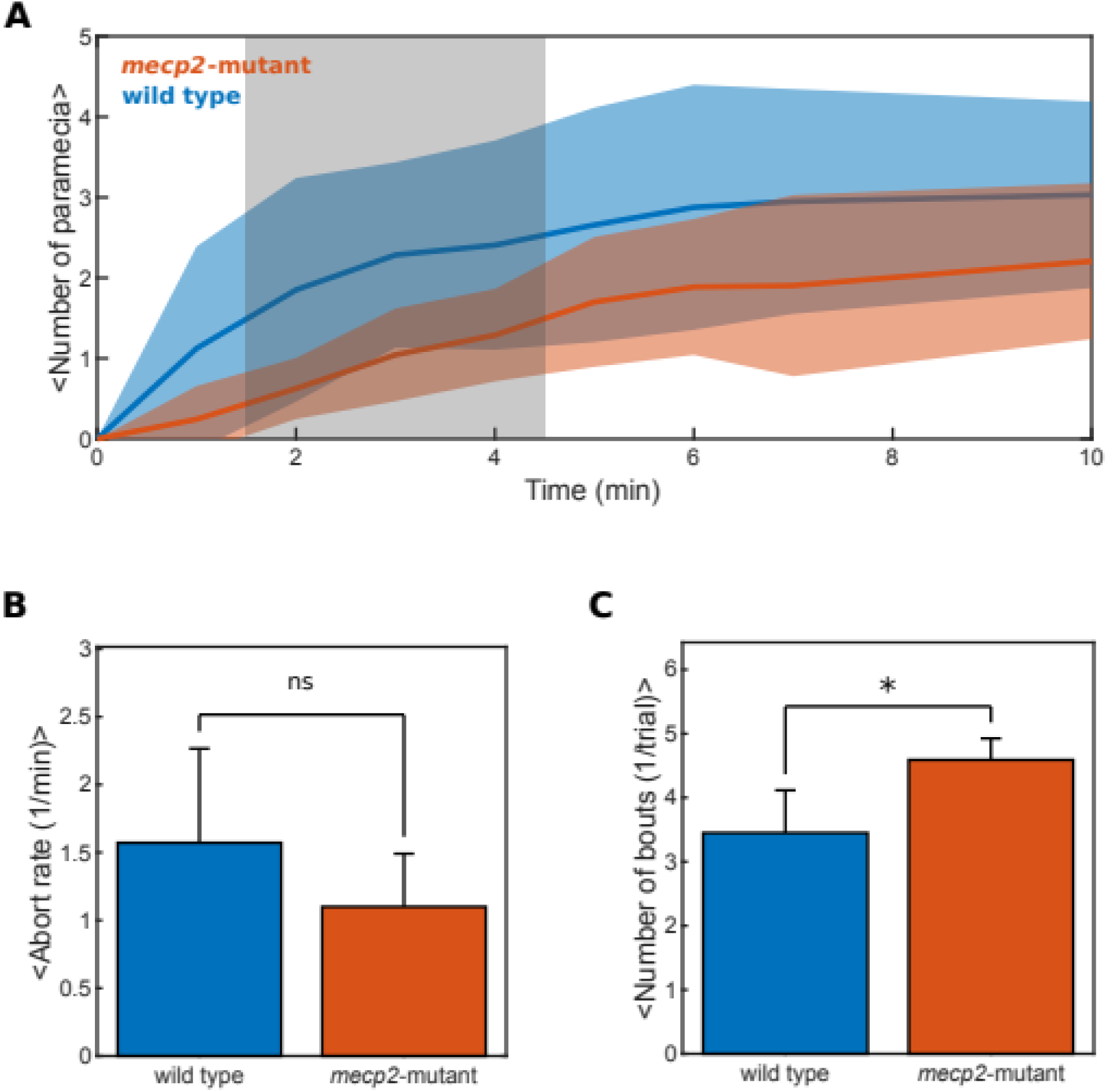
Prey-capture is mildly impaired in mecp2-mutant *larvae*. (A) The average number of paramecia eaten per larva for wild-type (blue, n=9) and *mecp2-*mutant (orange, n=9) larvae. Shaded area: standard deviation. Gray area: p<0.05, right-tailed Wilcoxon ranksum test. (B) The average abort rate for wild-type (blue, 1.57 ± 0.69, n=7) and *mecp2-*mutant larvae (orange, 1.10 ± 0.39, n=8), measured as the number of eye-convergence events where the larvae prematurely aborted prey capture behavior, per minute, during the first 5 minutes of recording. p=0.577, right-tailed Wilcoxon ranksum test. Error bars: standard error. (C) The average number of bouts per eye-convergence events for wild-type (blue, 3.45 ± 0.67, n=7) and *mecp2-*mutant larvae (orange, 4.59 ± 0.33, n=8), during the first 5 minutes of recording. p=0.045, left-tailed Wilcoxon ranksum test. Error bars: standard error.

To better understand this hypothesis, we recorded single larvae at 100 Hz. to characterize prey-capture kinematics. We found that on average, the rate of aborted prey-capture events (no capture strike at the end of the eye convergence episodes) was not significantly different between wild-type and *mecp2-*mutant larvae (wild type: 1.57 ± 0.69, n=7 versus *mecp2-*mutant: 1.10 ± 0.39, n=8; p=0.58 right-tailed Wilcoxon ranksum test, Fig. 4B). However, the total number of bouts per prey capture episodes was higher for *mecp2-*mutant than for wild-type larvae (wild type: 3.45 ± 0.67, n=7 versus *mecp2-*mutant: 4.59 ± 0.33, n=8; p=0.045 left-tailed Wilcoxon ranksum test, Fig. 4C and Supp. Movie2).

These results suggest that *mecp2-*mutant larvae are capable of capturing prey, but at a slower rate because they require a higher number of bout movements (corrections).

## DISCUSSION

Neuronal circuits with attractor-like dynamics are recurrently connected networks whose temporal dynamics settle into self-sustained stable activity patterns. These characteristics allow neuronal circuits to recall stored patterns from partial cues (i.e., pattern completion), or to discriminate and categorize its inputs (i.e, pattern separation). In zebrafish larvae, it was previously observed that the tectal circuit is organized according to functional neuronal assemblies ^18–21^ with attractor-like dynamics ^16^. Here, we investigated the functional role and the biological relevance of these tectal assemblies using mutant larvae (MeCP2Q63*, ^22^). This mutant showed a reduced local functional connectivity between tectal neurons, despite their normal lifespan, normal behavioral phenotypes, and a normal ratio between the number of excitatory (Vglut2a) and inhibitory (GAD1b) neurons. Therefore, the *mecp2* mutant represents a unique experimental model to study *in vivo* the functional role of the neuronal assemblies in the optic tectum.

In this study, we took advantage of the abnormal functional connectivity in the tectal circuit of the *mecp2*-mutant larvae, to assess the effect of the neuronal assemblies with attractor-like dynamics, on the visual responses and prey-capture behavior. We found that the *mecp2*-mutant larvae displayed an abnormal visual response. In comparison to wild-type larvae, the tectal response of the *mecp2*-mutant larvae to large visual stimuli of different contrasts had smaller amplitude, shorter duration, were less reliable and less sensitive. In contrast, the percentage of responsive neurons was larger for the *mecp2*-mutant larvae. These differences worsened the potential capacity of the tectal network of the *mecp2*-mutant to decode the contrast of the presented visual stimuli.

The presentation of small light-spots mimicking potential prey (*e.g*. paramecia) induced less reliable and smaller amplitude visual responses in the tectum of the *mecp2*-mutant larvae. In contrast to the neuronal responses to large visual stimuli, the percentage of responsive neurons induced by the small light-spot stimuli, was larger in wild-type larvae than in the *mecp2*-mutant larvae . These differences worsened the capacity of the tectal network of the *mecp2*-mutant to decode the azimuth position of the visual stimuli. Overall, these results suggest that the intrinsic tectal functional connectivity organized as neuronal assemblies with attractor-like dynamics increases the strength of the visual response (duration and integral), and improves visual sensitivity and response reliability. In addition, when visual stimuli are large, the neuronal assemblies restrain the number of visually induced neurons. This probably occurs through reciprocal inhibition between neighboring assemblies (winner-takes-all dynamics, ^16^). In contrast, when visual stimuli are small, the tectal intrinsic connectivity amplifies the visual response (increases the number of active neurons as in pattern complition). Behaviorally, the *mecp2*-mutant larvae also required a larger number of bouts before catching prey (more correction movements to capture the prey).

These properties correspond to those expected from attractor circuits capable of self-sustaining neuronal activity underlying eye fixation, memory and navigation ^2,9,10,12,31–33^, pattern discrimination involving decision making ^16,34,35^, and pattern completion supporting memory retrieval ^1–3^.

Our results suggest that the neuronal assemblies in the optic tectum ameliorates the visual spatial resolution and the signal-to-noise ratio, acting as a top-down virtual fovea, improving the detection sensitivity to prey-like stimuli with respect to visual stimuli of other sizes.

The hypothesis that abnormal brain connectivity underlies autism has been controversial ^36^. However, recent studies reconciled this controversy showing that different phenotypes of autism spectrum disorders (ASD) are associated with specific patterns of aberrant connectivity. For example, magnetic resonant studies in mice ^37^ and humans ^38^ showed that different gene mutations (*e.g.* Fmr1, CDLK5, Shank3, BTBR, MECP2) are associated with 4 different types of ASD phenotypes. Mutations leading to similar phenotypes showed a specific type of abnormal connectivity (hypo or hyper connectivity) in specific brain regions.

In humans, the single mutation of the *mecp2* gene is sufficient to induce Rett syndrome disease, with ASD symptoms ^39^. Our findings showed that the optic tectum of *mecp2*-mutant larvae displays hypo functional connectivity. This suggests that at least in the visual system, autistic-like phenotypes could be associated with hypo-connectivity altering the circuits’ attractor dynamics.

Furthermore, autistic patients show hyper-responsiveness and hyper-sensitivity to sensory stimuli ^40,41^. A previous study showed that *mecp2*-mutant zebrafish larvae display reduced thigmotaxis when freely swimming in an arena, potentially avoiding tactile stimulation from the wall ^22^. Here, we showed that the hypo-connectivity in *mecp2*-mutant larvae results in sensory responses involving a larger number of neurons. Thus, our results may shed light on the neuronal mechanism underlying hyper-sensitivity in autism.

## ACKNOWLEDGMENTS AND FUNDING

We would like to thank Thomas Tulinski and Lucie Daveau for conducting preliminary experiments. M.P. was supported by the Fondation pour la Recherche Medicale (FRM: FDT201904008327) and ENS Lyon, T.P. was supported by L’Association Française du syndrome de Rett, G.S. was supported by ERC CoG 726280.

## AUTHOR CONTRIBUTION

Conceptualization, T.P. and G.S.; methodology, M.P., E.H., T.P. and G.S.; software, M.P.; formal analysis, M.P., E.H., T.P., A.D. and E.M.; investigation, M.P., E.H., T.P., E.M., A.U-A., A.D, S.N.; resources, M.P., E.H., T.P., G.S., V.C., and S.N.; writing, M.P., E.H., T.P. and G.S.; supervision, G.S.; project administration, G.S.; funding acquisition, G.S, M.P., T.P..

## MATERIALS AND CORRESPONDENCE

Correspondence and materials requests should be addressed to Germán Sumbre

## METHODS

### Fish lines

All experimental procedures were approved by the comité d’éthique en expérimentation animale n◦005. Reference number APAFIS#27495-2020100614519712 v14. 2.2. All experiments were performed using zebrafish larvae from 5 to 7 days post-fertilization (dpf), expressing pan-neuronally the GCaMP3 ^16^ or GCaMP5g ^42^ indicator on a nacre (mitfa -/-) background ^43^, or on a nacre mecp2^Q63*/Q63^* mutant. The embryos were collected and raised at 28°C in 0.5x E3 embryo medium (E3 in mM: 5 NaCl, 0.17 KCl, 0.33 CaCl2, 0.33 MgCl2 pH 7.2). Larvae were kept under 14/10 h on/off light cycles and fed after 5 dpf with Paramecia.

### Transgenic zebrafish larvae

For the spontaneous activity and large visual stimuli experiments, we used a transgenic HuC:GCaMP3 transgenic zebrafish line ^44^. To create the line, we built the tol2 HuC:GCaMP3 vector using successive ligations of a 3.2 kb fragment of the zebrafish HuC (elav3) promoter ^45^ (gift from HC Park, Kyungpook National University, Korea) and then ligations of the GCaMP3 calcium probe ^46^ (gift from L. Looger, Howard Hughes Medical Institute, Ashburn, Virginia, USA) into pT2KXIG in (from K. Kawakami, National Institute of Genetics, Shizuoka, Japan). The HuC promoter drives the expression of an RNA-binding protein involved in neuronal differentiation. The 3.2 kb proximal region encompasses the translation start site from 2771 base pairs of the 5’-upstream sequence up to +383/+385. This fragment in zebrafish has been shown to be sufficient to target specifically and efficiently all differentiated neurons ^45^.

We injected 20 ng of the DNA plasmid and 25 ng of transposase RNA (generated from pCS-TP plasmid, K. Kawakami) to one-cell stage Nacre zebrafish embryos. The injected embryos were raised to adulthood and then crossed individually with Nacre fish to obtain F1 embryos. We screened these embryos and selected those with the strongest level of transgene expression. The selected embryos were raised to adulthood and crossed between each other to obtain the homozygous HuC:GCaMP3^GS5^ line (ens100Tg at ZFIN).

For the small light-spot visual stimuli (prey-like stimuli), we used 5-6 dpf Tg(HuC:GCaMP5G)^ens102Tg^ zebrafish larvae. To generate this line, we first built a tol2 HuC:GCaMP5G vector by insertion of a 3.2 kb fragment of the zebrafish HuC (elav3) pan-neuronal promoter ^45^. Next, the genetically encoded Ca^2+^ indicator GCaMP5G ^47^ was inserted into pT2KXIG (from K. Kawakami). Then, we injected one-cell-stage nacre zebrafish embryos with 10 ng of the plasmid DNA and 25 ng of transposase RNA which was generated from pCS-TP plasmid (K. Kawakami). The injected embryos reached adulthood and were crossed individually with nacre fish to obtain F1 embryos. Next, we screened and then selected the F1 embryos according to their level of transgene expression. Finally, the embryos with the highest expression were raised to adulthood and incrossed to obtain the homozygous HuC:GcaMP5G(^ens102Tg^)

### The MeCP2^Q63*/Q63^* mutant larvae

Zebrafish *mecp2^Q^*^63*/Q63^* mutation was generated through *N*-ethyl-*N*-nitrosourea (ENU)-mutagenesis and selected by TILLING ^22^. As ENU-mutagenesis generates random mutations throughout the genome, heterozygote *mecp2-*mutant fish were outcrossed several times in the AB background to remove off-target mutations. Mutant fish were identified by PCR using DNA extracted from fin clip (the primers 5′-AAAGGAAAGGCATGATGTGG-3′ and 5′-GTATCGCCAACCTTTTGGAA-3′ flank the position of the mutation), followed by sequencing.

HuC:GCaMP3, MeCP2*^Q^*^63*/Q63^* larvae and HuC:GCaMP5, MeCP2*^Q^*^63*/Q63^* were generated by crossing

HuC:GCaMP3 and HuC:GCaMP5 fish with homozygous MeCP2*^Q^*^63*/Q63^* fish to obtain F1 embryos. F1 fish were then incrossed, and their offspring were screened for green fluorescence and sequenced to keep only MeCP2*^Q^*^63*/Q63^* homozygous individuals.

### Quantification of excitatory and inhibitory neurons

#### Immunofluorescence

Zebrafish larvae (6 dpf) were fixed for 2 hours at 4°C in 4% paraformaldehyde in PBS, washed three times in PBS + 0.5% Triton, blocked one hour in PBS + Triton 0.5% + DMSO 1% + Goat serum 10%. For the excitatory neurons, the primary antibody used was a chicken anti-RFP (AB3528, Merck Millipore) diluted 1:300 in blocking buffer for overnight incubation. As the secondary antibody, we used the goat anti-chicken coupled to Alexa Fluor 568 (A11041, Life technologies), diluted 1:500 in blocking buffer for overnight incubation. For the inhibitory neurons, the primary antibody used was a rabbit anti-GFP antibody (TP401, Torrey pines biolabs) diluted 1:1000 in blocking buffer for overnight incubation. For the secondary antibody, we used the goat anti-rabbit coupled to Alexa Fluor 488 (A11034, Life technologies), diluted 1:500 in blocking buffer for overnight incubation. After the third wash, the larvae were stained with 10 μg/mL of 4, 6-diamidino-2-phenylindole (DAPI, D9542, sigma) in PBS + triton 1% + DMSO 1% + Tween 0.1% overnight, mounted in 2% low melting agarose, and imaged using confocal microscopy. Images were collected using a Leica 25× water immersion objective (HC FLUOTAR L, NA 0.95, working distance 2.5 mm) with an upright Leica laser-scanning confocal TCS SP5 II (Leica Microsystems, Germany) equipped with photomultiplier tubes (PMT), a motorized stage, standard- and high-resolution Z-focus, and laser lines (405, 458, 476, 488, 514, 561, and 633 nm). Images were acquired using LAS AF software.

#### Segmentation and cell-type detection

In order to perform cell segmentation and cell-type detection, we used a custom made program in Matlab. The three channels recorded (DAPI, gad1b and Vglut2a) were cropped using a mask to delineate the area of the optic tectum. To identify the cell bodies of the neurons, we performed a 2D cross-correlation between the images and the template of a neuron (2D Gaussian of the size and dispersion matching those of a common zebrafish neuron, ∼ 5 μm). The outcome was binarized using Otsu’s method, which finds a threshold that minimizes the intra-class variance of the thresholded black and white pixels ^48^. Regions larger or smaller than 5 μm were discarded. Cells whose binary mask for the Vglut2a and gad1b channels had less than 45% overlap with the binary masks for the DAPI channel were discarded.

The average intensity of each channel (Vglut2a and/or gad1b) was computed as the average value for the ROI segmented using the DAPI channel. We classified cells as Vglut2a or gad1b positive if the average intensity was 50% higher in one of the channels compared to the other.

### Calcium Imaging

#### Two-photon recordings

We used an imaging set-up based on a MOM system (Sutter) with a 25x NA 1.05 Olympus objective and a Mai Tai DeepSee Ti:sapphire laser tuned at 920 nm. Three different Semrock filters were used: an FF705 dichroic (objective dichroic), an AFF01-680 short path (IR Blocker) and an FF01 520/70 band-pass filter. The PMT was a H1070 (GaAsP) from Hamamatsu. The emission signal was pre-amplified with a SR-570 (Standford Research Systems). We acquired tiff images using ScanImage ^49^ at 3.91 Hz or 1.95 Hz at a resolution of 256 x 256 pixels.

#### Spontaneous activity

We recorded 10 *mecp2-*mutant and 9 wild-type larvae expressing GCaMP3 at 5 dpf during 35 minutes. The larvae were left a few minutes to adapt to the experimental environment before starting the recordings.

#### Wide-field looming stimuli

We recorded 10 *mecp2-*mutant and 7 wild-type larvae expressing GCaMP3 at 6 dpf. The larvae were subjected to 10 increasing light intensity stimuli (from 0.1x to 1x projector capability), with a 7s inter-stimulus interval. Each stimulus lasted 1s and consisted of a fast looming (150ms for full expansion) of 40° angle of the visual field centered at 30° angle with respect to the head of the larva. The stimuli were repeated 15 times. High contrast values (0.9 and 1x) resulted in a saturated response in both *mecp2*-mutant and wild-type larvae. Consequently, we only analyzed contrast values from 0.1x to 0.8x.

#### Light-spot stimuli

We recorded 9 *mecp2-*mutant and 7 wild-type larvae expressing GCaMP5 at 6 or 7 dpf. Small (5°) light-spot stimuli were presented at four different azimuth position contralateral to the recorded tectum (15°, 35°, 40° and 50° with respect to the anteroposterior axis where 0° is in front of the larva’s head). Each visual stimulus was presented for 1s with a 7s inter-stimulus interval, and repeated 20 times. Stimuli were shuffled and presented in a pseudo-random order. To maximize tectal responses, a small vertical jitter of 10° was added along the elevation axis (vertical speed of 20° per second). The imaging plane of the tectal circuit was chosen according to the maximal response to the presented stimuli.

### Behavior Recordings

#### Paramecia count

To compare the prey-capture kinematics between wild-type and *mecp2-*mutant larvae, we recorded 9 experiments with wild types and 9 experiments with *mecp2-*mutant larvae, and counted the number of paramecia remaining at different time points during the experiment (30 min).

In each trial, 6 or 7 dpf larvae were placed inside a small rectangular chamber (20x20 mm). Their behavior was recorded at 100 Hz. from above with an infrared camera (Baumer TXG02), and illuminated from below with an infrared LED (850 nm). For wild-type larvae, each trial was composed of 5 larvae, and for *mecp2-*mutant larvae 7 trials were composed of 5 larvae and 2 of 4 larvae. At the beginning of the experiment, we introduced paramecia in the well and recorded for 30 minutes. We introduced on average 24.8 ± 8.8 paramecia. Transparent agarose was placed along the sides of the chamber to facilitate the detection of both larvae and paramecia.

We measured the number of paramecia in the chamber every minute from 0 to 7 min, and every 5 min from 10 to 30 min. Each measurement was done by recording images from 4 consecutive seconds and taking the difference with the first image, to see the displacement of objects (larvae and paramecia), and counting by hand the paramecia. Each time point reported in the data is the mean of 3 measurements.

#### Tracking

To analyze in more details the kinematics of prey-capture behavior, we used the same setup with a single larva for each experiment and tracked the position of the larva as well as the convergence of the eyes and shape of the tail ^50^.

We identified prey-capture events as periods of high convergence of the eyes (using a bivariate Gaussian mixture model with two components for the angle of the long axis of each eye relative to the anteroposterior axis).

### Data analysis

#### Preprocessing of calcium imaging

The pre-preprocessing of the calcium imaging data was done according to Romano et al. ^16,17^. Briefly, for each animal, each frame of the recording was registered to a reference image using a cross-correlation approach to cancel out any potential drift in the X-Y axes. Movements artifacts due to struggle movements of the larva in the agarose were identified as large transient X-Y drifts in the cross-correlation analysis, and the corresponding frames removed from the analysis. Neurons were automatically segmented using a watershed-based algorithm, and the resulting ROIs were manually curated. All the pixels of each ROI were averaged to compute the relative changes in fluorescence (ΔF/F) with respect to the baseline, computed as the 8^th^ percentile of a sliding window of 30s ^51^. Significant calcium events (SCE) were extracted using a Bayesian inference method ^16^ relying on the estimation of the baseline noise for each neuron, where SCE(i,j)=0 if there is no significant calcium event at time *i* for neuron *j*, and SCE(i,j)=1 if there is a significant calcium event.

#### Correlation analysis and Jensen-Shannon distance

To compute the correlations between neurons, we used the original value of ΔF/F where significant calcium events were identified, and replaced the value of ΔF/F with 0 everywhere else (baseline equal to zero). This method makes the data more sparse, and may remove correlations linked to seconds-long fluctuations that may have remained after the computation of the baseline (which acts as a high pass filter).

To compute the average probability density function (PDF) for correlations in wild-type and *mecp2-* mutant larvae, we computed the histogram of correlation values for each larva in regularly spaced (0.001) correlation bins between -1 and 1. We then averaged those histograms together within each population.

To assess the differences in the level of correlations between wild-type and *mecp2-*mutant larvae, it is important to estimate the amount of correlations that would result from the basal level of activity of each cell alone. To do so we shuffled circularly the time series of each neuron by a different, random amount of time. The shuffling effectively destroyed the synchrony between neurons present in the original dataset without disturbing the firing rate or the distribution of inter-event-times.

This process was repeated 500 times to obtain the null-model for correlations. To compare the distributions of correlations between wild-type and *mecp2-*mutant larvae, we quantified the distance between the null-model distributions and the distribution of correlations in our dataset. For this purpose, we used the Jensen-Shannon distance for each larva, which is a measure of distance between histograms based on the Kullback-Leibler divergence ^52^.

Finally, to investigate the spatial structure of correlations in the optic tectum of wild-type and *mecp2-*mutant larvae, we averaged the correlations between pairs of neurons. The correlations were computed using regularly spaced (15 *μm*) distance bins between 0 and 240 *μm* to obtain the distribution of correlations against distance.

#### Selection of responsive cells

In order to select neurons that were responsive to the visual stimuli, we compared the neuronal activity in a time window of 3 s before and 3 s after the onset of each stimulus.

To assess significance, we used a non-parametric random sampling procedure to draw samples from those time windows with replacement (bootstrapping). For each bootstrap sample (1000 repetitions total), we computed the difference in the proportion of significantly active time frames (significant calcium event) after the onset vs. before the onset of each stimulus.

A stimulus trial was deemed responsive if the 95^th^ percentile of the bootstrapped distribution was strictly positive (significantly higher proportion of active frames after than before stimulus onset). A cell was deemed responsive to a stimulus type if it was active for more than X trials of this stimulus. The value of X was determined so that applying the bootstrap procedure to a sham stimulus onset (arbitrarily affected to a period of spontaneous activity) would not yield more than 5% of responsive cells on average over all larvae (X=2 trials out of 15 repetitions in Fig. 2, and X=1 trials out of 20 repetitions in Fig. 3).

Finally, a cell was deemed generally responsive if it was responsive to at least one stimulus type.

#### Maximum likelihood decoder

For the decoding, we used the same approach as described in ^53^. Briefly, we computed the average of the ΔF/F on a time window of 3 s after each stimulus onset as a measure of the neuronal response. A leave-one-out procedure was used for training: for a given stimulus type, each stimulus trial was predicted using the remaining (n_trials - 1) observations of all stimulus types. Continuous probability estimates were obtained by smoothing the training data with a Gaussian kernel function. The conditional probability of the observed neuronal response given the stimulus type for the population, was computed as the product of the conditional probabilities of all neurons (we assumed that the responses of neurons were independent).

#### Decoding the neuronal response consisted in finding the stimulus type that maximizes this conditional probability

The performance of the decoder was defined as the proportion of trials that were properly classified over all stimulus types (this is also called accuracy). The classification performance was computed using the fluorescence signals of 10 randomly chosen ensembles of n neurons and then averaged over ensembles and larvae.

To assess statistical significance of the classification performance, we calculated the probability of getting correct classifications by chance. This was given by the binomial distribution B(n,p), where p is the probability of getting a correct classification by chance (p=1/(number of stimuli)), and n is the number of tests (number of stimuli x number of repetitions). Significant decoding was reached when the decoding performance exceeded the 95^th^ percentile of the binomial distribution.

#### Stimulus response properties

In order to characterize the neuronal response to the different visual stimuli, we computed the integral of the response, peak response probability, proportion of responsive cells, and the response duration. To compute these properties, we first averaged the ΔF/F values over trials and responsive cells for each stimulus (Fig. 2), or overall cells that responded to at least one stimulus among all stimuli (Fig. 3), to get the quantity <ΔF/F>[trials,cells].

We applied the same procedure to the significant calcium events (SCE, see preprocessing of the data), to get the quantity <SCE>[trials,cells], which represents the frame-by-frame probability of activation across trials averaged over cells.

The integral of the response was defined as the sum of <ΔF/F>[trials,cells] on a time window of 6 s after the onset of the stimulus.

The peak response probability was defined as the maximum value of <SCE>[trials,cells] during a time window of 6 s after the onset of the stimulus.

The proportion of responsive cells was defined as the number of responsive cells for each stimulus divided by the total number of cells recorded for each larva (Fig. 2), or the number of cells responsive to at least one stimulus divided by the total number of cells recorded (Fig. 3).

Finally, the response duration was quantified as the full width at half-maximum value of <ΔF/F>[trials,cells] on a time window of 6 s after the onset of the stimulus.

#### Statistical inference, data and code availability

Non-parametric statistical tests were performed when the normality assumption could not be assumed. No statistical methods were used to determine sample sizes in advance, but sample sizes are similar to those reported in other studies in the field. Multiple testing correction was not applied when looking for the location of statistical differences along curves after two-way ANOVAs.

Details about the statistical tests (one or two-tailed), number of samples used, and type of error bars (standard deviation vs standard error of the mean) are defined in the text and figure legends.

#### Raster plots

Raster plots were filtered using a high-pass filter with a frequency 0.03Hz to remove slow fluctuations in the time series.

## DATA AVAILABILITY

Data can be made available on request.

## CODE AVAILABILITY

Data analysis was performed using custom scripts written in MATLAB. The code can be made available on request. Pre-processing code for the calcium imaging data (https://github.com/zebrain-lab/Toolbox-Romano-et-al) and behavioral tracking (https://github.com/zebrain-lab/Tracking) can be found on github.

---

**Supplemental figure 1, related to figure 1:**
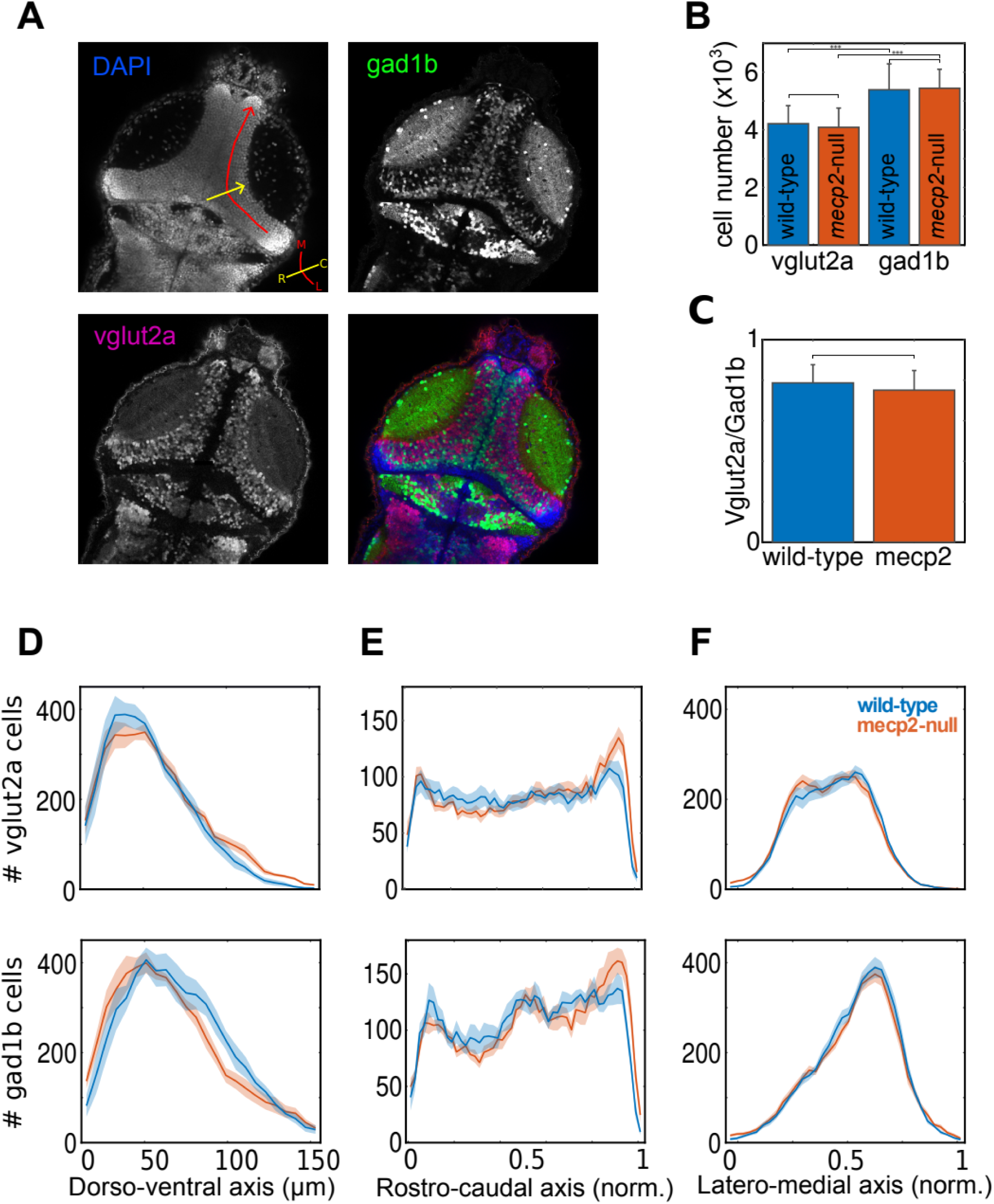
Cell type identity in the tectum of wild-type and mecp2-mutant larvae. (A) An optical section of a 6 dpf larva immunostained for DAPI (blue, top-left, nuclei of all neurons), gad1b (green, top-right, GABAergic neurons), and vglut2a (magenta, bottom-left, glutamatergic neurons). The bottom-right panel shows the superposition of the other 3 panels. Axes: R: rostral; C: caudal; M: medial; L: lateral. (B) The number of GABAergic (gad1b) and glutamatergic (vglut2a) neurons in the optic tectum of wild-type (blue, n=8) and *mecp2-*mutant (orange, n=8) larvae. No significant differences were found (wild type gad1b: 4.21±0.63, wild type vglut2a: 5.39±0.90, p= 6.6^-24^, GLME, *mecp2-*mutant gad1b: 4.09±0.67, *mecp2-*mutant vglut2a: 5.44±0.66, p= 2.4^-27^, GLME, wild type vglut2a vs. *mecp2-*mutant vglut2a: p=0.92, GLME, wild type gad1b vs. *mecp2-*mutant gad1b: p=0.86, GLME). (C) The inhibition-excitation ratio in wild type (blue, n=8) and *mecp2-*mutant (orange, n=8) larvae. No significant differences were found (wild type: 0.79±0.09, *mecp2-*mutant: 0.75±0.10, p = 0.43, GLME). (D-F) The number of glutamatergic (vglut2a, top) and GABAergic (gad1b, bottom) cells for wild-type (blue) and *mecp2-*mutant (orange) larvae, distributed along the dorso-ventral axis (vglut2a: p =0.69, gad1b: p=06, GLME) (D), along the rostro-caudal axis (vglut2a: p=0.7, gad1b: 0.83, GLME) (E), and the latero-medial axis (vglut2a: p=0.78, gad1b: p=0.69, GLME)(F), in wild-type (blue) and *mecp2-*mutant (orange) larvae. See axes on (A).

**Supplemental movie 1 related to figure 3**:

Example of tectal activity in response to small light dot stimuli in a wild-type (left) and *mecp2*-mutant larva (right). Neuronal activity was averaged across trials for each stimulus location, represented by circles of different colors (blue: 15°, cyan: 35°, yellow: 40°, and red: 50°). The spatial location was calculated using a 2D kernel density estimation method. The small colored circles represent the center of mass of the neurons responsive to the visual stimulus (density peak). The dotted colored lines indicate the spatial extent of the response (level curve of the density at half-maximum). The period of the visual stimulation and its azimuth position are indicated by the filled colored circles on the top-left corner. The intensity of the neuronal activity was normalized to the 99^th^ percentile of the ΔF/F values for each stimulus azimuth separately, and color coded in red for each neuron. The video represents 1.75x of its real speed. Note the differences between wild-type and *mecp2*-mutant larvae with respect to the center of mass of the responses, and the spatial extent of the response. In contrast to the *mecp2*-mutant larva, the wild-type larva, displayed well separated centers of mass of the visual responses, revealing the retinotipic map of the optic-tectum. Also, the visual responses in the tectum of the wild-type larva involved a larger number of neurons in a more spatially constrained area.

**Supplemental movie 2, related to figure 4**:

Example of a prey capture event for a wild-type (left) and *mecp2-*mutant larva (right). Red circles: paramecia, green line: skeleton of the larva’s tail, black line: antero-posterior axis of the larva, white line: mediol-ateral axis of the larva, red line: major axis of the right eye, cyan line: major axis of the left eye. Note that the *mecp2*-mutant overshoots when performing the j-turns, requiring more j-turns before capturing the paramecium.

